# Expression of elongase- and desaturase-encoding genes shapes the cuticular hydrocarbon profiles of honey bees

**DOI:** 10.1101/2024.07.22.604606

**Authors:** Daniel Sebastián Rodríguez-León, Thomas Schmitt, María Alice Pinto, Markus Thamm, Ricarda Scheiner

## Abstract

Most terrestrial insects have a layer of cuticular hydrocarbons (CHCs) protecting them from desiccation and mediating chemical communication. CHC composition is regulated by the expression of genes coding for enzymes in the biosynthetic pathway of hydrocarbons. The diversity and expression of these enzymes determine the abundance and richness of compounds in the CHC profile of an insect. For example, elongases are enzymes that lengthen the hydrocarbon chain, while desaturases introduce double bonds in it. CHC profiles of honey bees (*Apis mellifera*) vary among castes, task groups, and subspecies. This makes *A. mellifera* an excellent model to study the molecular mechanism underlying CHC biosynthesis. Here, we examined the expression of specific elongase- and desaturase-encoding genes and correlated gene expression with CHC composition in bees from two different task groups of two highly divergent *A. mellifera* subspecies: *A. m. carnica* and *A. m. iberiensis*. We show that in *A. mellifera*, the specificity of desaturases and elongases shapes the CHC profiles of different task groups. Our results shed light on the genetic basis for the task-specific CHC composition differences in social hymenopterans and lay the ground for further studies aiming to unravel the genetic underpinning of CHC biosynthesis. Moreover, these results underline the importance of investigating different subspecies of *A. mellifera* to better understand the mechanisms driving CHC composition.

## Introduction

Cuticular hydrocarbons (CHCs) protect insects from water loss and mediate inter- and intraspecific communication (Blomquist et al. 1987; Gibbs 1998; Howard and Blomquist 2005). The composition of the CHC layer determines its biological function (Blomquist et al. 1987; Howard and Blomquist 2005). For example, its ability to act as a desiccation barrier depends on the aggregation of its constituent hydrocarbons (Gibbs 1995; Gibbs 2002; Menzel et al. 2019). The n-alkanes aggregate denser than methyl-branched or unsaturated hydrocarbons, due to stronger van der Waals forces (Gibbs and Pomonis 1995; Gibbs 1998). The aggregation of hydrocarbons also increases with their chain length (Gibbs and Pomonis 1995). The denser the aggregation of the hydrocarbons, the more waterproof is the CHC layer (Gibbs 1995; Gibbs 2002; Howard and Blomquist 2005). But a dense aggregation of hydrocarbons also increases the viscosity of the CHC layer, thereby limiting hydrocarbon diffusion and interindividual CHC exchanges and constraining the function of CHCs as communication cues (Gibbs 1995; Gibbs and Pomonis 1995; Gibbs 2002; Menzel et al. 2019).

CHCs are primarily synthesized by oenocytes, specialized secretory cells associated with the fat bodies of insects (Wigglesworth 1933; Schal et al. 1998; Makki et al. 2014; Moris et al. 2023). The CHC biosynthetic pathway is closely related to fatty acid metabolism. It is initiated with the synthesis of fatty acyl-coenzyme A (acyl-CoA), from the binding of malonyl-CoA units to acetyl-CoA (Howard and Blomquist 2005; Blomquist and Bagnères 2010). Along this pathway, several enzymes modify the fatty acyl-CoA to ultimately produce the different CHCs (Chung and Carroll 2015; Holze et al. 2021). Some of these enzymes contribute to the elongation of the hydrocarbon chain (elongases) (Cinti et al. 1992; Denic and Weissman 2007), while others introduce unsaturations (double bonds) into the hydrocarbon chain (desaturases) (Dallerac et al. 2000; Takahashi et al. 2001). The abundance and richness of CHC compounds on the cuticle of an insect are determined by the diversity and expression of the enzymes participating in this biosynthetic pathway (Juárez et al. 1992; Reed et al. 1995; Gu et al. 1997; Qiu et al. 2012). The expression of desaturase-encoding genes influences the diversity and abundance of unsaturated hydrocarbons (e.g., alkenes and alkadienes), while the expression of elongase-encoding genes influences the diversity and abundance of compounds with specific chain lengths. However, little is known about how the expression of genes encoding for enzymes of the same type (e.g. different elongases and desaturases) contribute to the variability of insect CHC composition.

A great example of how gene expression differences can lead to differential phenotypes can be found in social insects (i.e. hymenopterans), making them excellent models to study the relationship between gene expression and complex phenotypes (Smith et al. 2008; Hunt et al. 2010; Hunt et al. 2013). In honey bees (*Apis mellifera*), the CHC profiles of the workers differ according to their age, task-performance, and subspecies (Kather et al. 2011; Vernier et al. 2019; Rodríguez-León et al. 2024). The CHC profiles of nurse bees are characterized by a higher abundance of unsaturated hydrocarbons and longer chain lengths compared to forager bees (Kather et al. 2011; Rodríguez-León et al. 2024). These two social task groups typically differ in age. Young workers typically care for the brood and the queen (nurse bees), while the oldest workers leave the colony to collect pollen and nectar (forager bees) (Huang and Robinson 1996). Because the expression of specific elongase- and desaturase-encoding genes varies with age (Vernier et al. 2019), the differences in the CHC composition between nurses and foragers assumedly respond to a developmental regulatory process.

The differences in the CHC profiles of nurse and foragers bees are present in different *A. mellifera* subspecies (Rodríguez-León et al. 2024). *A. mellifera* subspecies have divergent evolutionary histories, leading to both genetic and phenotypic differences (Ruttner et al. 1978; Ruttner 1988; Garnery et al. 1992; Dogantzis et al. 2021). Different subspecies have evolved distinct CHC profiles, exhibiting both quantitative and qualitative differences in their composition (Rodríguez-León et al. 2024). This remarkable variability of CHC profiles makes *A. mellifera* an excellent model for studying the molecular mechanisms underlying CHC biosynthesis. A recent study elucidated a clear role of genes encoding for specific elongases and desaturases in the CHC biosynthesis of honey bee workers (Moris et al. 2023). A knock down of these genes resulted in compositional changes in CHC profiles, aligning with the expected function of the respective enzyme type. However, the role of desaturase and elongase gene expression in shaping CHC profiles has remained unclear.

Here, we correlated the expression of elongase- and desaturase-encoding genes in nurse and forager honey bees of two highly divergent subspecies, *A. m. carnica* and *A. m. iberiensis*, with their CHC profile composition. We hypothesized that the shift in CHC composition during the transition from nursing to foraging is driven by changes in the expression of specific CHC biosynthesis-related genes. These changes in gene expression should be preserved in both subspecies. We also hypothesized that the subspecies-specific CHC compositions of honey bee workers is influenced by differences in the expression of specific CHC biosynthesis-related genes between subspecies.

## Materials and methods

We performed a common garden experiment, mantaining queen-right colonies of *Apis mellifera carnica* and *Apis mellifera iberiensis* in the departmental apiary of the University of Würzburg (Germany). Although these subspecies are both native to Europe, they belong to the most evolutionarily divergent lineages of *A. mellifera* (Wallberg et al. 2014). *A. m. carnica* belongs to the Western European C-lineage, whereas *A. m. iberiensis* belongs to the Eastern European M-lineage (Ruttner 1988). *A. m. carnica* is native to the Central and Southeastern parts of Europe, but it has been introduced worldwide, and it is now the dominant subspecies in Germany (Maul and Hähnle 1994). *A. m. iberiensis* is native to the Iberian Peninsula, where it experienced historical secondary contact with the African A lineage (Ruttner 1988; De la Rúa et al. 2009; Han et al. 2012; Chávez-Galarza et al. 2015).

The colonies were assembled on June 2, 2021, using artificial swarms obtained from *A. m. carnica* colonies and mated queens from both subspecies. *A. m. carnica* queens were obtained from local German beekeepers, and *A. m. iberiensis* queens were obtained from local beekeepers in Portugal.

Honey bee workers were collected between the 22nd and 23rd of September 2021, between 9 a.m. and noon. Ten nurse bees and ten forager bees were taken from two colonies per subspecies, for a total of 80 worker bees. The 40 nurse bees were piked while poking their heads into a brood cell for at least 10 seconds. The 40 forager bees were piked while landing at the hive entrance. Only bees with conspicuous pollen loads were collected. The workers were collected into individual vials and transported to the lab on ice, where they were killed by storing them at −80 °C. The collected bees were preserved at −80 °C until the extraction of their CHCs.

### CHC composition analysis

The bees were defrosted and immersed in hexane for only two minutes to extract their CHCs and avoid RNA degradation. Immediately after the extraction with hexane, the bees were dissected to remove their guts and separate the abdomen from the rest of the body. The dissected abdomens were preserved separately in 963 µl of RNA-later, stored at 4 °C for 48 hours, and then at −20 °C until their RNA was extracted. The CHC extracts were stored at −20 °C.

The CHC extracts were analyzed via gas chromatography/mass spectrometry (GC/MS) by injecting 1µl of each extract into an Agilent 7890A Series GC System coupled to an Agilent 5975C Mass Selective Detector (Agilent Technologies, Waldbronn, Germany) operating in electron impact ionization mode at 70 eV and a source temperature of 230 °C. The split/splitless injector was operated in splitless mode for 1 minute at 300 °C. Separation of compounds was performed on a J&W DB-5 fused silica capillary column (30 m × 0.25 mm ID, df = 0.25 µm, J&W, Folsom, California, USA) with a temperature program starting at 60 °C and increasing by 5 °C per minute to 300 °C, which was held for 10 minutes. Helium served as a carrier gas with a constant flow of 1 ml per minute.

The chromatograms were analyzed using the data analysis software package “MassHunter Workstation Software - Qualitative Analysis Navigator” (version B.08.00; Agilent Technologies, Inc. 2016). The areas of the peaks were determined by integration using the “Agile2” parameter-less integrator, with the peak filters set to 0% of the area of the largest peak. The integration results were aligned regarding the retention time (RT) of the peaks for all the samples of each colony and task-performance (e.g., nurse or forager bees) group. The differences in performance of the total ion counts were corrected based on n-alkane analytical standard solutions (04070-1ML and 04071-5ML, Sigma-Aldrich). The aligned group data sets were merged by the Kováts retention index (RI) and compound identity of the peaks. Compounds that represented less than 0.01% of the total ion count of a sample, and compounds detected in less than 50% of the samples within a group, were excluded from the analysis. Only the compounds identified as alkanes, alkenes, alkadienes, or methyl-branched alkanes were considered in the analysis. The different hydrocarbons in the extracts were identified based on their diagnostic ions and RIs. Double bond positions of monounsaturated hydrocarbons (alkenes) were identified by dimethyl disulfide derivatization (Carlson et al. 1989). The abundances of the compounds were quantified for each sample as the proportion (%) that the area of the corresponding peaks represented from the sum of the area of all the peaks included in the analysis.

Additionally, we calculated the relative abundance of both alkenes and alkadienes, as well as the mean chain length, in the CHC profile of each bee and compared them between tasks and between subspecies. The relative abundances of the alkenes and alkadienes (mono- and diunsaturated hydrocarbons, respectively) were calculated as the sum of the relative abundances of all the individual alkenes or alkadienes found in the CHC extract of a bee. The mean chain length corresponds to the weighted mean of the chain length of all the compounds found in the CHC profile of each bee, using the relative abundance of the compounds as their weights.

### Gene expression analysis

The GenUpTM total RNA kit (Biotechrabbit, Henningsdorf, Germany) was used to extract the total RNA from each of the dissected abdomens, following the manufacturer’s instructions. Along the RNA extraction procedure, a DNase I digestion step was performed by adding 30 µl DNase mix containing 30 U RNase-free DNase I (Lucigen Corporation, Middleton, USA) together with the corresponding buffer, after the binding of the RNA to the Mini Filter RNA. The total RNA concentration was determined photometrically, and the extracted RNA was diluted in DEPC-H_2_O to a concentration of 100 ng/µl and stored at −80 °C until the synthesis of the cDNA. A total of 400 ng of RNA was used for cDNA synthesis with the cDNA synthesis kit 331475S/L (Biozym, Hessisch Oldendorf, Germany). The resulting cDNA solution was diluted up to a total volume of 180 µl with DEPC-H_2_O, and stored at −20 °C.

The expression of two desaturase-encoding genes (*Des1* and *Des2*; Table S1) and two elongase-encoding genes (*Elo1* and *Elo2*; Table S1) was determined via quantitative Polymerase Chain Reaction (qPCR). These four genes have been shown to affect CHC biosynthesis (Moris et al. 2023). Individual cDNAs were analyzed in triplicate for each gene using the SYBR Green BlueMix qPCR kit (Biozym, Hessisch Oldendorf, Germany) and a Rotor-Gene Q (Qiagen, Hilden, Germany). The following cycling conditions were set up for every qPCR run: 2 minutes at 95 °C, 40 cycles with 5 seconds at 95 °C, and 30 seconds at 60 °C. A melting curve was recorded by increasing the temperature from 60 °C to 95 °C in steps of 1 °C every 5 seconds. Four qPCR runs were performed per gene, including the reference genes (*Rpl10* and *Rpl19*; Table S1), for a total of 16 qPCR runs. Each qPCR run was performed with five samples per task and subspecies group. All the samples of both subspecies within a qPCR run corresponded to the same hive, and the same combination of samples per hive per run was used for the four genes. The expression of the four genes was quantified relative to the reference genes, following the qBase algorithm (Hellemans et al. 2007). Cycle threshold (Ct) values of sample replicates that deviated by more than 2 from the median Ct value of the corresponding sample were considered technical outliers and discarded from the calculation of the relative expression of the corresponding gene.

### Data analysis

We compared the relative abundance of alkenes and alkadienes, as well as the mean chain length, in the CHC profile of the bees between tasks (nurse bees vs. forager bees) and subspecies (*A. m. carnica* vs. *A. m. iberiensis*). We fitted quantile regression models for the 50% quantile, considering both task and subspecies as independent variables (predictors). The interaction between task and subspecies was only considered for the model cases where it was significant (i.e., the relative abundance of the alkenes).

To evaluate the difference in gene expression of the four assessed genes between tasks and subspecies, we performed bootstrapped gamma generalized linear model (GLM), with a logarithmic link function, over 10,000 resamples. Inference was made with the bootstrapped marginal means, model coefficients (standardized effect sizes), and their 95% confidence intervals, which were calculated via the percentile intervals method. Due to the use of the logarithmic link function, the model coefficients were transformed by exponentiation. Thus, they represent the proportional difference in gene expression to the reference group (task: nurse bees; subspecies: *A. m. carnica*).

We evaluated the correlation of the relative expression of both desaturase-encoding genes to the relative abundance of both alkenes and alkadienes, as well as to the relative abundance of every unsaturated hydrocarbon in the CHC profiles of the bees. Similarly, for the elongase-encoding genes, we evaluated the correlation of their relative expression to the mean chain length of the CHCs of the bees, as well as to the relative abundance of every compound in the CHC profiles of the bees. All correlations were calculated using Spearman’s correlation index. In addition, we performed a bi-dimensional Non-Metric Multidimensional Scaling (NMDS) using the Bray-Curtis dissimilarity index to assess the dissimilarity between samples, regarding the composition of their CHC profiles. Then, for each of the four assessed genes, we plotted on the NMDS the compounds with a positive (> 0.4) or negative (< −0.4) correlation towards the expression of the corresponding gene.

The data analysis was performed using R v4.4.0 (R Core Team 2024) and RStudio IDE v2024.4.1.748 (Posit Team 2024). Data wrangling and plotting operations were done using the packages tidyverse v2.0.0 (Wickham et al. 2019), ggtext v0.1.2 (Wilke and Wiernik 2022), gghalves v0.1.4 (Tiedemann 2022), ragg v1.3.2 (Pedersen and Shemanarev 2023), and patchwork v1.2.0 (Pedersen 2023). The processing of the integrated chromatograms’ data (i.e., RT wise alignment, RI calculation, abundance correction, peak filtering, and calculation of the relative abundance of compounds) was done using the packages analyzeGC v0.2.1 (Rodríguez-Leon 2023a) and GCalignR v1.0.7 (Ottensmann et al. 2018). The relative expression of the genes of interest was calculated using the package easyqpcr2 v0.1.0 (Rodríguez-Leon 2023b). The bootstrapped GLMs and quantile regression models were fit with the packages tidymodels v1.2.0 (Kuhn and Wickham 2020) and quantreg v5.98 (Koenker 2023) respectively. The marginal means were estimated using the package emmeans v1.10.3 (Lenth 2023). The NMDS was performed using the package vegan v2.6.6.1 (Oksanen et al. 2022). Spearman’s correlation indexes and their 95% confidence intervals were calculated using the package confintr v1.0.2 (Mayer 2023).

## Results

### CHC composition differs between bees performing different tasks and between subspecies

*A. m. carnica* nurse bees exhibited a higher abundance of both alkenes and alkadienes, as well as longer chain length hydrocarbons than the forager bees (Figures 1 and S1). In *A. m. iberiensis*, the nurse bees displayed a higher abundance of alkadienes and longer chain length hydrocarbons than the forager bees. Subspecies did not affect the abundance of alkadienes (Figures 1B and S1B). However, the CHC profile of *A. m. carnica* contained a higher abundance of alkenes and longer chain length hydrocarbons than that of *A. m. iberiensis*, in both the nurse and forager bees (Figures 1C and S1C).

**Figure 1:**
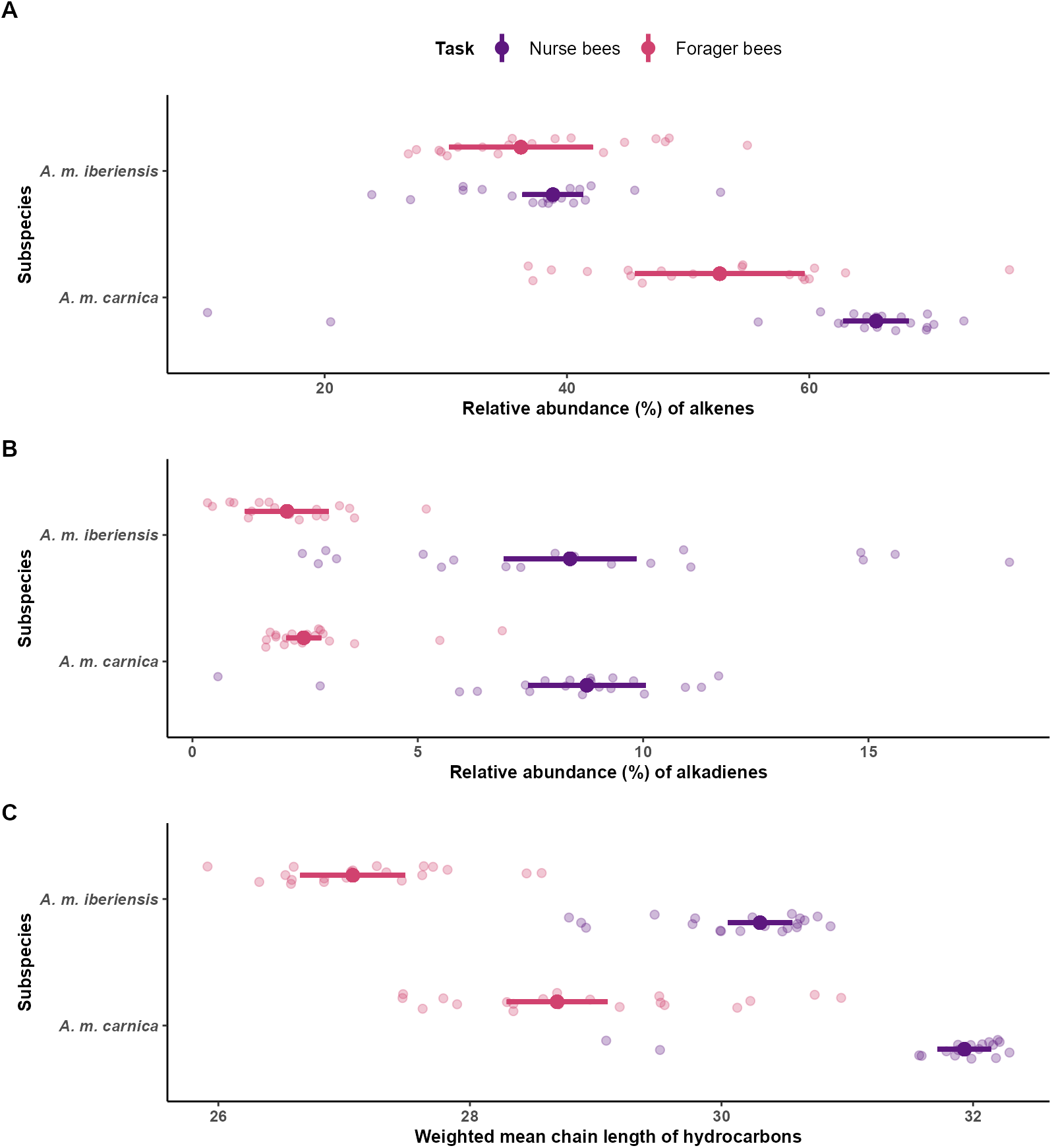
Task and subspecies-related differences in the CHC compositions of honey bee workers (nurse and forager bees) of two subspecies (*A. m. carnica* and *A. m. iberiensis*). The figure is divided into three plots, each corresponding to the results of a quantile regression analysis on the task- and subspecies-related differences in a compositional trait of the CHC profiles of the bees. **A)** Relative abundance of mono-unsaturated hydrocarbons (alkenes). **B)** Relative abundance of di-unsaturated CHCs (alkadienes). **C)** Mean chain length of the hydrocarbons. Point intervals represent the model prediction for the median and its 95% confidence interval. The raw data is visualized with transparent points, which correspond to the measured amount in the CHC profile of every bee.

### Nurse and forager bees differ in the expression of desaturase- and elongase-encoding genes

The expression of the two desaturase-encoding genes differed between nurse and forager bees in both subspecies (Figures 2 and S2). Forager bees expressed *Des1* 0.295 times as much as nurse bees (95% CI: 0.240, 0.363). Conversely, the expression of *Des2* in the forager bees was 2.209 times higher than in the nurse bees (95% CI: 1.722, 2.819).

**Figure 2:**
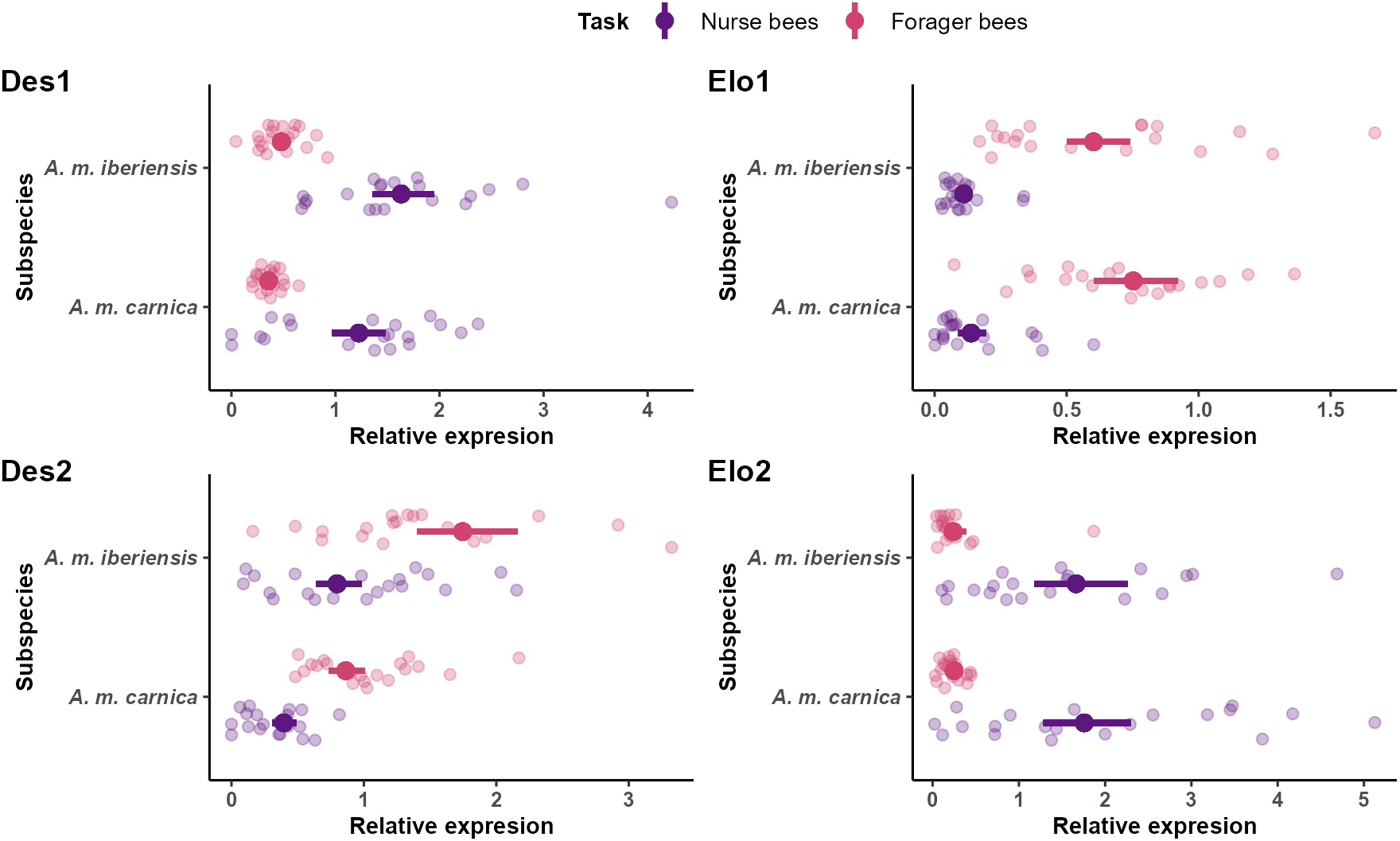
Relative expression of CHC biosynthesis-related genes in honey bee workers (nurse and forager bees) of two subspecies (*A. m. carnica* and *A. m. iberiensis*) genes. The figure is divided into four plots, each corresponding to a gene. Point intervals represent the median of the bootstrapped GLM prediction for the mean relative expression of the genes and its 95% confidence interval. The raw data is visualized with transparent points, which correspond to the relative gene expression value that was measured for every honey bee worker.

Nurse and forager bees also differed in the expression of both elongase-encoding genes in both subspecies (Figures 2 and S2). *Elo1* was expressed 5.586 times more by the forager bees than the nurse bees (95% CI: 4.053, 7.774). On the other hand, the expression of *Elo2* in the foragers was 0.145 times that of the nurse bees (95% CI: 0.095, 0.214).

### Subspecies differ in the expression of desaturasebut not elongase-encoding genes

*A. m. iberiensis* elicited a higher expression of both desaturase-encoding genes than *A. m. carnica*, for both nurse and forager bees (Figures 2 and S2). The expression of *Des1* was 1.346 times higher in *A. m. iberiensis* than in *A. m. carnica* (95% CI: 1.092, 1.658). Moreover, the expression of *Des2* was 2.039 times higher in *A. m. iberiensis* than in *A. m. carnica* (95% CI: 1.594, 2.593). Subspecies did not differ in the expression of *Elo1* (95% CI: 0.589, 1.135) or *Elo2* (95% CI: 0.639, 1.426). This pattern was observed for both nurse and forager bees (Figures 2 and S2).

### Correlation between gene expression and CHC composition

Bees with a higher relative expression of *Des1* exhibited a higher abundance of alkadienes in their CHC profiles (Figure 3A). In contrast, the relative expression of *Des2* was negatively correlated with the abundance of both alkenes and alkadienes.

**Figure 3:**
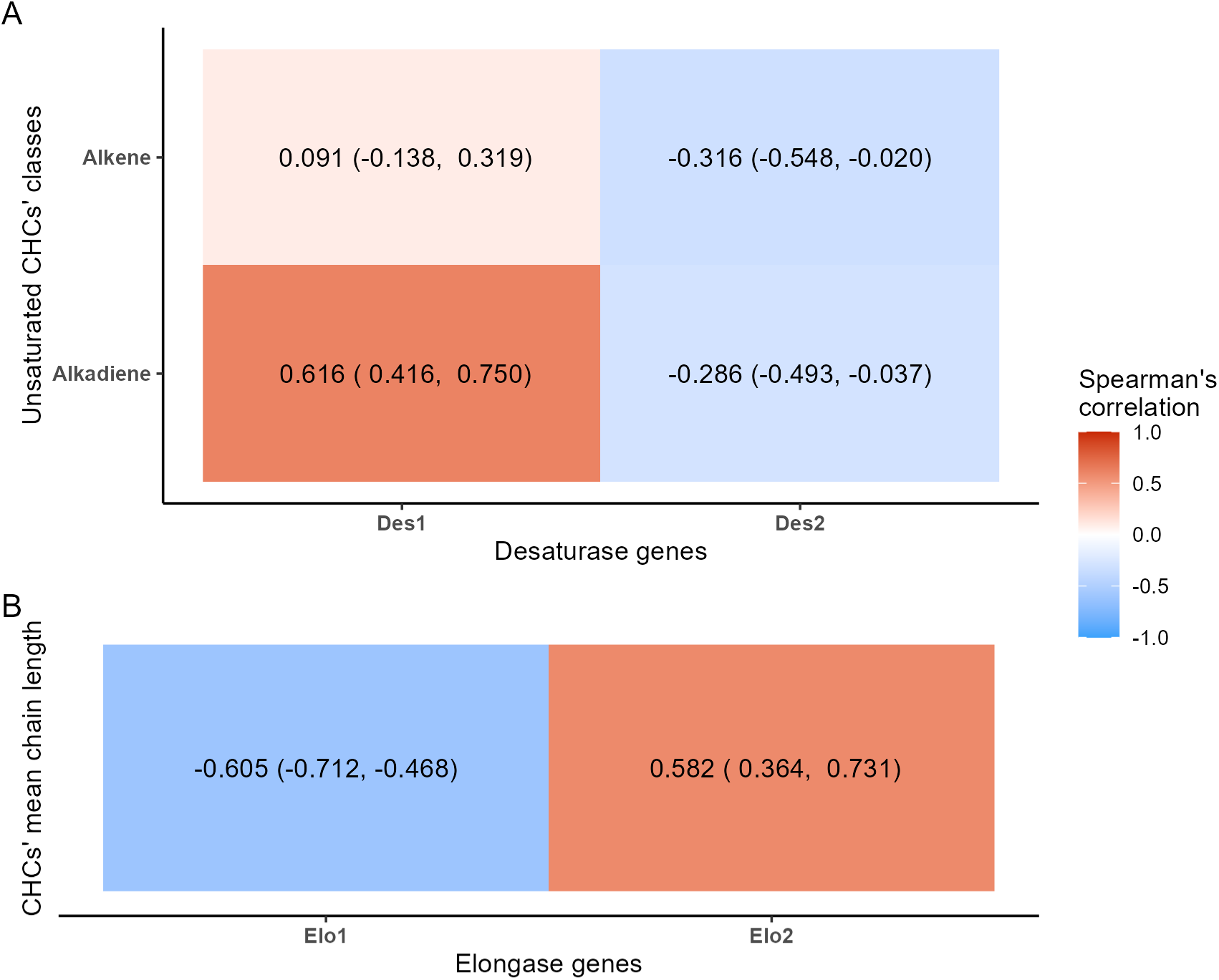
Correlation of the expression of CHC biosynthesis-related genes and CHC composition in honey bee workers (nurse and forager bees) of two subspecies (*A. m. carnica* and *A. m. iberiensis*). **A)** Correlation between the relative expression of both desaturase-encoding genes and the relative abundance of alkenes and alkadienes (%). **B)** Correlation between the relative expression of both elongase-encoding genes and the weighted mean chain length of the CHCs. The estimated Spearman’s correlation index is shown besides its 95% confidence interval (lower limit, upper limit).

Bees with a higher expression of *Des1* tended to have a higher abundance of unsaturated hydrocarbons with chain lengths over 29 carbon atoms in their CHC profiles (Figure 4A). Bees with a higher expression of *Des2* displayed higher abundances of unsaturated compounds with up to 29 carbon atoms in their CHC profiles. The compounds that positively correlated with the expression of *Des1* were more abundant in the CHC profile of nurse bees, while those negatively correlating with the expression of *Des1* were more abundant in the CHC profiles of forager bees (Figure 4B). *A. m. iberiensis* nurse bees had a higher abundance of the hydrocarbons that positively correlated with the expression of *Des1* than the nurse bees of *A. m. carnica*. In contrast, forager bees of *A. m. iberiensis* exhibited a higher abundance of the hydrocarbons that negatively correlated with the expression of *Des1* than their *A. m. carnica* counterparts. In the case of *Des2*, forager bees displayed a higher abundance of the hydrocarbons that were positively correlated with its expression. Nurse bees, in turn, presented a higher abundance of the hydrocarbons that negatively correlated with the expression of *Des2*. *A. m. iberiensis* forager bees presented a higher abundance of the hydrocarbons that positively correlated with the expression of *Des2* than the forager bees of *A. m. carnica*. The nurses of *A. m. carnica*, on the contrary, displayed a higher abundance of the hydrocarbons that negatively correlated with the expression of *Des2* than their *A. m. iberiensis* counterparts.

**Figure 4:**
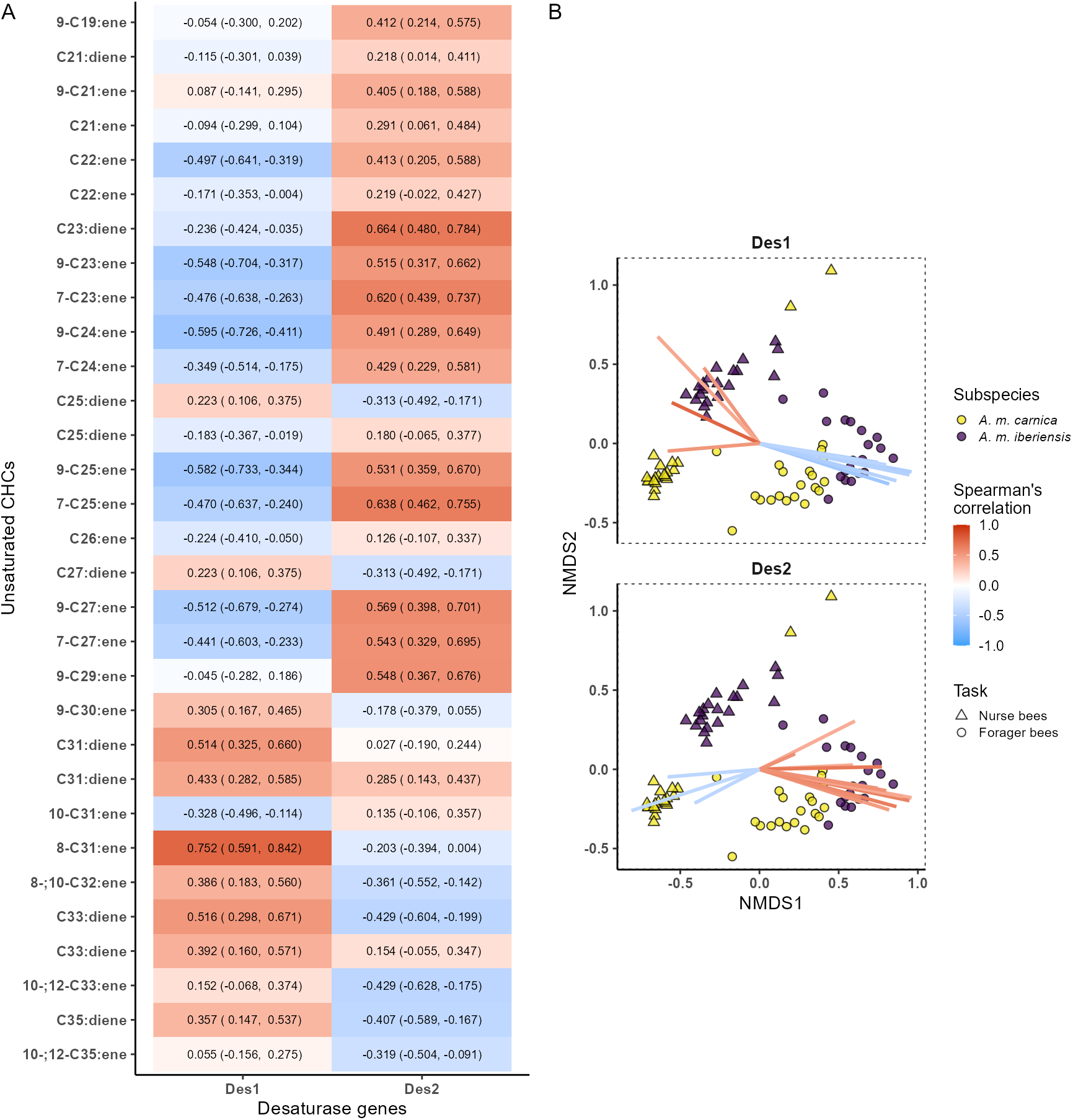
Correlation between the expression of desaturase-encoding genes and the CHC composition of honey bee workers (nurse and forager bees) of two subspecies (*A. m. carnica* and *A. m. iberiensis*). **A)** Correlation between the relative expression of both desaturase-encoding genes and the relative abundance (%) of hydrocarbons. The hydrocarbons are sorted from shorter chain length (top) to longer chain length (bottom). The estimated Spearman’s correlation index is shown besides its 95% confidence interval (lower limit, upper limit). **B)** Bi-dimensional Non-metric Multidimensional Scaling (NMDS) analysis on a dissimilarity matrix of the CHC profiles of honey bee workers. Ordination stress: 0.083. The plot is divided into facets, each depicting the same NMDS but only illustrating the coordinates of the CHCs correlated with the expression of a specific desaturase gene. The colored lines illustrate the coordinates of the CHCs. The estimated correlation between hydrocarbons’ abundance (%) and the expression of the corresponding gene is indicated by the color of the line (red: positive. blue: negative). Only hydrocarbons with a correlation higher than −0.4 (negative) or 0.4 (positive) are shown.

Bees with a higher expression of *Elo1* tended to have shorter compounds in their CHC profiles (Figure 3B). More specifically, the expression of *Elo1* correlated positively with the abundance of compounds with chain lengths between 22 and 27 carbon atoms and negatively with the abundance of longer compounds (Figure 5A). On the contrary, bees with a higher expression of *Elo2* had the tendency to exhibit CHC profiles with longer compounds (Figure 3B). The higher the expression of *Elo2*, the higher was the abundance of the longer compounds (≥ 27 carbon atoms), in particular of those with chain lengths over 29 carbon atoms. In turn, compounds with chain lengths shorter than 27 carbon atoms tended to be less abundant in the CHC profiles of the bees with a higher expression of *Elo2*.

**Figure 5:**
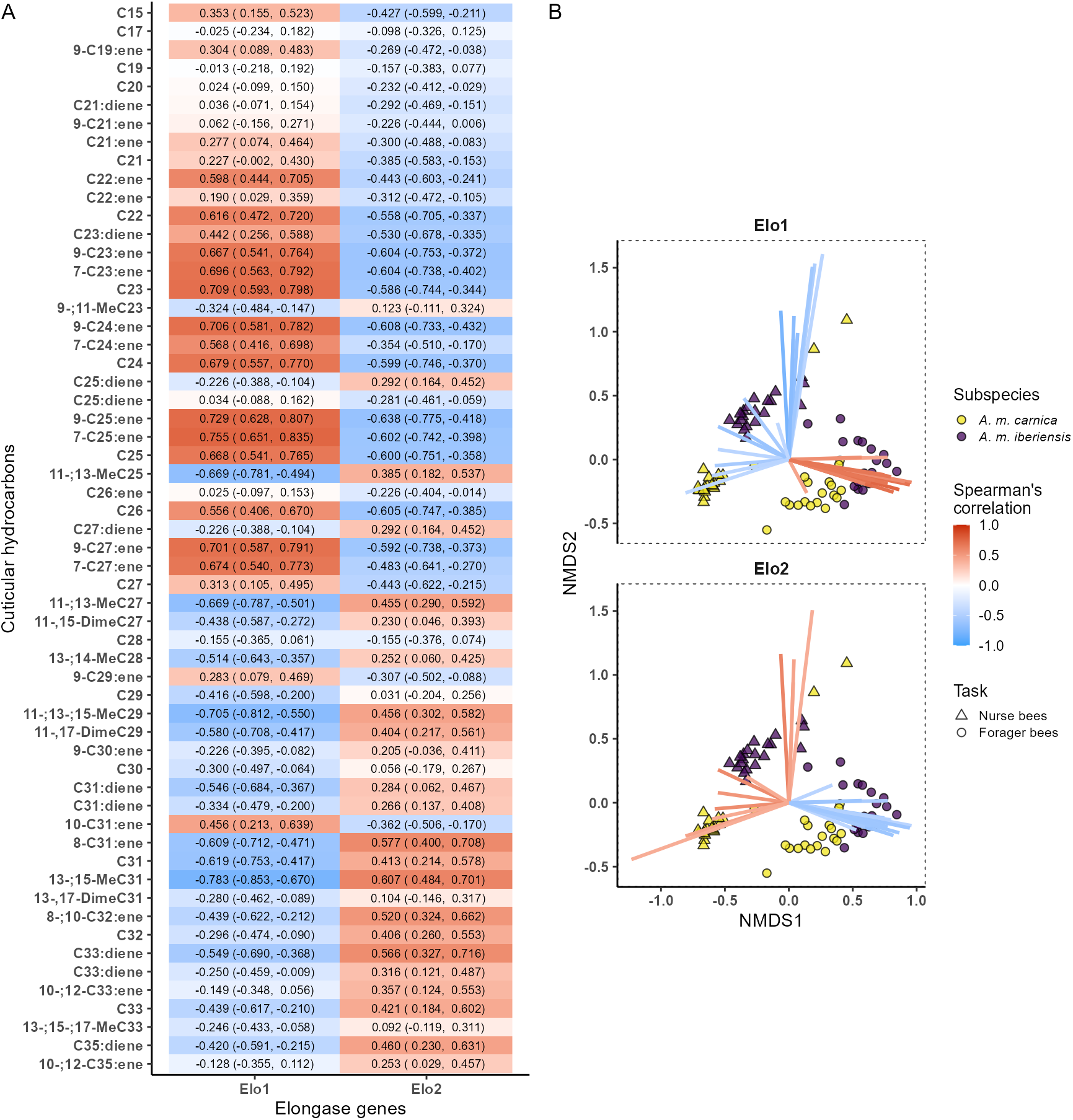
Correlation between the expression of elongase encoding genes and the CHC compositions of honey bee workers (nurse and forager bees) of two subspecies (*A. m. carnica* and *A. m. iberiensis*). **A)** Correlation between the relative expression of both elongase genes and the relative abundance (%) of hydrocarbons. The hydrocarbons are sorted, top-to-bottom, from shorter to longer chain length. The estimated Spearman’s correlation index is shown besides its 95% confidence interval (lower limit, upper limit). **B)** Bi-dimensional Non-metric Multidimensional Scaling (NMDS) analysis on a dissimilarity matrix of the CHC profiles of honey bee workers. Ordination stress: 0.083. The plot is divided into facets, each depicting the same NMDS but only illustrating the coordinates of the CHCs correlated with the expression of a specific elongase gene. The colored lines illustrate the coordinates of the CHCs. The estimated correlation between hydrocarbons’ abundance (%) and the expression of the corresponding gene is indicated by the color of the line (red: positive. blue: negative). Only hydrocarbons with a correlation higher than −0.4 (negative) or 0.4 (positive) are shown.

The hydrocarbons with the strongest correlation with the expression of both elongase-enconding genes were highly associated with the social task (Figure 5B). *Elo1* correlated positively with the hydrocarbons that were more abundant in the CHC profiles of forager bees and negatively with those that were more abundant in the CHC profiles of nurse bees. *A. m. iberiensis* forager bees displayed a higher abundance of the compounds that were positively correlated with the expression of *Elo1* compared to *A. m. carnica* forager bees. This difference between subspecies was less evident among nurse bees. However, most of the hydrocarbons that correlated negatively with the expression of *Elo1* were more abundant in the *A. m. iberiensis* nurse bees than in their *A. m. carnica* counterparts.

The compounds that correlated positively with the expression of *Elo2* were more abundant in nurse bees compared to forager bees. On the contrary, forager bees displayed a higher abundance of the compounds that correlated negatively with the expression of *Elo2* than nurse bees. Here, the differences between the two subspecies seem to only be evident between the forager bees, with *A. m. iberiensis* displaying a higher abundance of the compounds that were negatively correlated with the expression of *Elo2* than *A. m. carnica*.

## Discussion

### Nurse and forager bees differ in their CHC compositions

Our data revealed a different CHC composition between nurse and forager bees. Nurse bees typically exhibited compounds with longer chain lengths and a higher abundance of unsaturated compounds in their CHC profile compared to forager bees. This difference was stereotypically consistent for two evolutionary divergent *A. mellifera* subspecies. Furthermore, it suggests that the CHC layer of forager bees protects them better against desiccation compared to nurse bees (Florian Menzel et al. 2017; F. Menzel et al. 2017; Menzel et al. 2018). A similar difference has been described for the workers inside and outside the nest of other social insects (Wagner et al. 1998; Wagner et al. 2001; Martin and Drijfhout 2009). This is thought to respond to the higher desiccation pressure faced by outside workers (e.g., forager bees) compared to those inside the nest (e.g., nurse bees), due to their exposure to weather extremes (e.g., temperature and humidity) (Wagner et al. 1998; Wagner et al. 2001; Martin and Drijfhout 2009; Kather et al. 2011; Rodríguez-León et al. 2024). Moreover, this difference in CHC composition between workers inside and outside the nest might serve as cues to inform other workers about the task they perform (Greene and Gordon 2003; Martin and Drijfhout 2009).

Honey bee workers exhibit an age-dependent division of labor, transitioning from nursing to foraging as they age. This transition is accompanied by a series of physiological changes (M. Elekonich et al. 2001; Johnson 2008; Siegel et al. 2013; Reim and Scheiner 2014; Scheiner et al. 2014; Scheiner et al. 2017; Schilcher and Scheiner 2023). The observed differences in the CHC composition of nurse and forager bees could therefore be influenced by shifts in the expression of CHC biosynthesis-related genes that occur along their task transition.

### Underlying mechanism of the task-related CHC composition shift in honey bees

The four examined genes (*Des1*, *Des2*, *Elo1*, and *Elo2*) strongly differed in their expression between nurse and forager bees. *Des1* and *Elo2* were up-regulated in nurse bees and down-regulated in forager bees, while the opposite pattern was observed for *Des2* and *Elo1*. Honey bees division of labor is flexible and responds to the needs of the colony, allowing individual bees to transition from nursing to foraging at different ages (Winston and Fergusson 1985; Huang and Robinson 1992; Huang and Robinson 1996; Johnson and Frost 2012). In fact, by artificially manipulating the colony, forager bees can be induced to revert to nurse bees, both behaviorally and physiologically (Robinson et al. 1992). Therefore, it is plausible that the observed differences in the expression of CHC biosynthesis-related genes between nurse and forager bees respond to regulatory changes associated with task transition.

We found that the difference in the expression of CHC biosynthesis-related genes between nurse and forager bees correlates with their different CHC compositions. Specifically, the expression of *Des1* and *Elo2* correlated positively with the abundance of compounds that were characteristic of nurse bees. The expression of *Des2* and *Elo1*, on the other hand, correlated positively with the abundance of compounds that were characteristic of forager bees. These findings suggest a yet unknown fine-tuning mechanism involving the up- and down-regulation of these genes, influencing the shift in the CHC composition of honey bee workers during their transition from nursing to foraging. Intriguingly, this transcriptional mechanism appears to be conserved across evolutionary lineages, as is suggested by the consistent correlation between task-specific CHC profiles and the expression of these genes in both *A. mellifera* subspecies.

### A. m. carnica and A. m. iberiensis differ in CHC composition

*A. m. carnica* displayed longer compounds and a higher abundance of unsaturated compounds in their CHC profiles than *A. m. iberiensis*. These results suggest that *A. m. iberiensis* is better adapted to a higher drought stress than *A. m. carnica* (Florian Menzel et al. 2017; F. Menzel et al. 2017; Menzel et al. 2018). However, differences in CHC composition between *A. mellifera* subspecies have been suggested to respond to their divergent evolutionary histories and genetic drift, as they do not seem to consistently correlate with the climatic differences (i.e., temperature and precipitation) between their native distributions (Rodríguez-León et al. 2024). Therefore, the adaptive difference in CHC composition we observed between *A. m. carnica* and *A. m. iberiensis* is likely specific to this particular comparison. These two subspecies belong to different *A. mellifera* evolutionary lineages and their natural ranges encompass very different environmental conditions (Ruttner 1988; De la Rúa et al. 2009; Han et al. 2012; Chávez-Galarza et al. 2015). As our study was a common garden experiment, we attribute the observed differences in the CHC composition between *A. m. carnica* and *A. m. iberiensis* to innate factors, leading us to expect an innate difference in expression of CHC biosynthesis-related genes between the two subspecies.

### *A. mellifera* subspecies differences highlight CHC biosynthesis complexity

*A. m. iberiensis* exhibited a higher expression of both desaturase-encoding genes (*Des1* and *Des2*) than *A. m. carnica*. In contrast, the expression of both elongase-encoding genes (*Elo1* and *Elo2*) did not differ between subspecies. These findings suggest that the expression of *Des1* and *Des2* between *A. mellifera* subspecies would respond to their evolutionary divergence, but not the expression of *Elo1* and *Elo2*. Since we performed a common garden experiment, the similarity in the expression of *Elo1* and *Elo2* by the bees of both subspecies could indicate an influence of the environment on the expression of these two elongase-encoding genes. However, our results do not discard the possibility that the expression these genes responds to intrinsic factors that are not related to the subspecies.

Nevertheless, among the genes assessed here, only the expression of *Des2* suggests a plausible mechanism underlying the differences in the CHC composition between *A. m. carnica* and *A. m. iberiensis*. The expression of this gene is positively correlated with the abundance of hydrocarbons with chain lengths of up to 29 carbon atoms. In consequence, the higher expression of *Des2* in *A. m. iberiensis* compared to *A. m. carnica* could contribute to the shorter mean chain length of hydrocarbons in the CHC profile of *A. m. iberiensis* compared to *A. m. carnica*. The expression of *Des1*, on the other hand, correlated with the abundance of alkadienes, a feature not differing between subspecies. Similarly, while the CHC profile of *A. m. carnica* exhibited longer hydrocarbons compared to *A. m. iberiensis*, the expression of both elongase-encoding genes (*Elo1* and *Elo2*) did not differ between subspecies. These discrepancies between the subspecific differences in CHC composition and the expression of the four studied genes evidence the complexity of CHC biosynthesis. Although several genes have been proposed to participate in CHC biosynthesis in honey bees, only a few have been mechanistically confirmed through targeted knock-down experiments to directly influence CHC composition in worker bees (Falcón et al. 2014; Vernier et al. 2019; Moris et al. 2023). It is thus plausible that the observed subspecific differences in CHC composition stem from the activity of the enzymes encoded by other CHC biosynthesis-related genes.

### Differential roles of specific genes in CHC biosynthesis

Our findings suggest that *Des1* and *Des2* play distinct roles in CHC biosynthesis. The expression of *Des1* correlated positively with the total relative abundance of alkadienes in the CHC profile of the bees, while the expression of *Des2* correlated negatively with the abundance of both alkenes and alkadienes. We consider that this difference could stem from the specificities of the enzymes encoded by *Des1* and *Des2*. Different desaturases have been found to vary in their substrate specificities and, thereby, in how they influence CHC composition (Dallerac et al. 2000; Takahashi et al. 2001; Holze et al. 2021). In our experiments, the expression of the two desaturase-encoding genes correlated with the abundance of different unsaturated hydrocarbons, supporting the above hypothesis. Moreover, the suggested difference in specificities between the enzymes encoded by these two genes explains the negative correlation between the expression of *Des2* and the abundance of both alkenes and alkadienes. In contrats to *Des1*, the expression of *Des2* expression was positively correlated with the abundance of unsaturated hydrocarbons with chain lengths of up to 29 carbon atoms. These compounds were the least abundant in the CHC profiles of honey bee workers, considering the mean chain length of the hydrocarbons in all analyzed groups (nurse and forager bees of both *A. m. carnica* and *A. m. iberiensis*) (Figure 1C). These compounds therefore only contributed little to the total abundance of alkenes and alkadienes.

In the case of the elongase-encoding genes, we found that *Elo1* expression negatively correlated with the mean chain length of the CHCs, unlike *Elo2*. While this *Elo1* pattern seems counterintuitive, as elongases catalyze the elongation of the hydrocarbon chain (Cinti et al. 1992; Jakobsson et al. 2006; Chung and Carroll 2015), this result can be explained by a differential specificity of the elongases encoded by the two genes. Elongases have been found to generally catalyze the biosynthesis of hydrocarbons with chain lengths of over 20 carbon atoms. Thus, the difference in specificity among elongases would explain the diversity of chain lengths in the CHC profiles of insects (Denic and Weissman 2007; Blomquist and Bagnères 2010; Holze et al. 2021). The expression of *Elo1* and *Elo2* correlated with the abundance of compounds within different chain length ranges, suggesting that both genes influence the synthesis of hydrocarbons with different chain lengths.

The diversity and regulation of the expression of CHC biosynthesis-related genes are thought to influence the richness and abundance of compounds in the CHC profile of an insect (Juárez et al. 1992; Reed et al. 1995; Gu et al. 1997; Howard and Blomquist 2005; Blomquist and Bagnères 2010; Qiu et al. 2012; Chung and Carroll 2015; Holze et al. 2021). However, current knowledge of the differential roles of enzymes of the same type (e.g., specific desaturases) shaping CHC composition is limited. Our data shed light on this topic by revealing correlations between the expression of specific desaturase- and elongase-encoding genes and CHC compositions in *A. mellifera* workers.

### Key Insights and Future Directions

Our study reveals conserved correlations between CHC composition and expression of specific CHC biosynthesis-related genes in honey bees. We provide evidence for a differential specificity among individual desaturases and elongases and propose that differences in the expression of their encoding genes contribute to shaping the CHC profiles of *A. mellifera* workers during their task transition from nursing to foraging. These results lay the ground for further studies aiming to unravel the genetic underpinning of CHC biosynthesis, in particular regarding task-specific CHC composition differences among social hymenopterans. We suggest further investigating *A. mellifera* subspecies to better understand the mechanisms underlying the CHCs diversity of honey bees. The complexity of the genetic basis underlying the difference in CHC composition among *A. mellifera* subspecies could reveal exciting insight into the molecular basis of CHC biosynthesis.

## Acknowledgements

We are grateful to Karin Möller and Nicole Martinez for their technical assistance during the experiments. We also would like to acknowledge Martin Gabel for his invaluable guidance in beekeeping. A special gratitude is extended to Douglas B. Sponsler and Diego Andrés Benítez Duarte for the stimulating conversations and insightful comments.

## Data availability

The data and code used for this study are not included in this manuscript, but will be made publicly available as supplementary material for the corresponding paper at the time of publication.

## Supplementary material

**Table S1:**
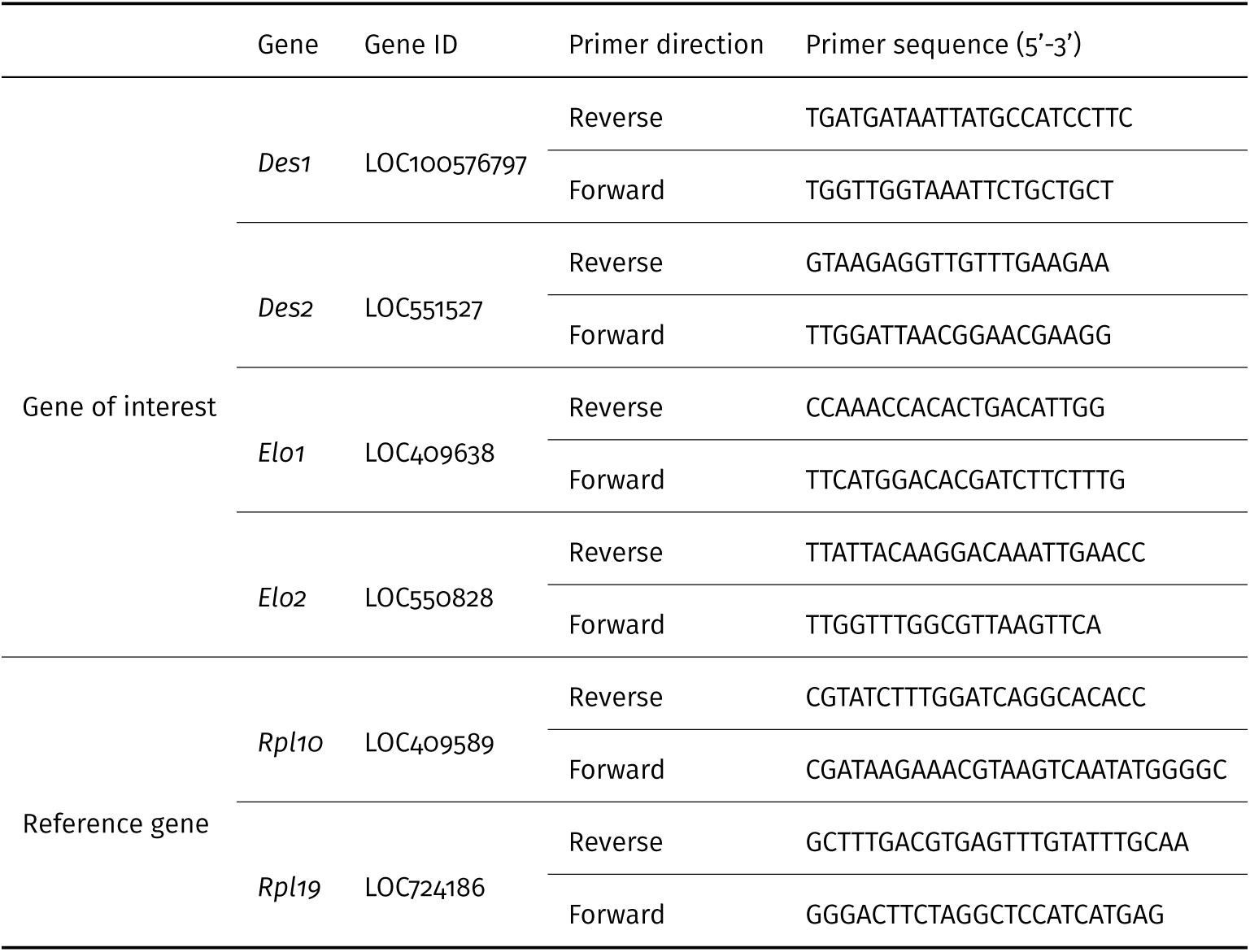
Oligonucleotide primers used in this study.

**Figure S1:**
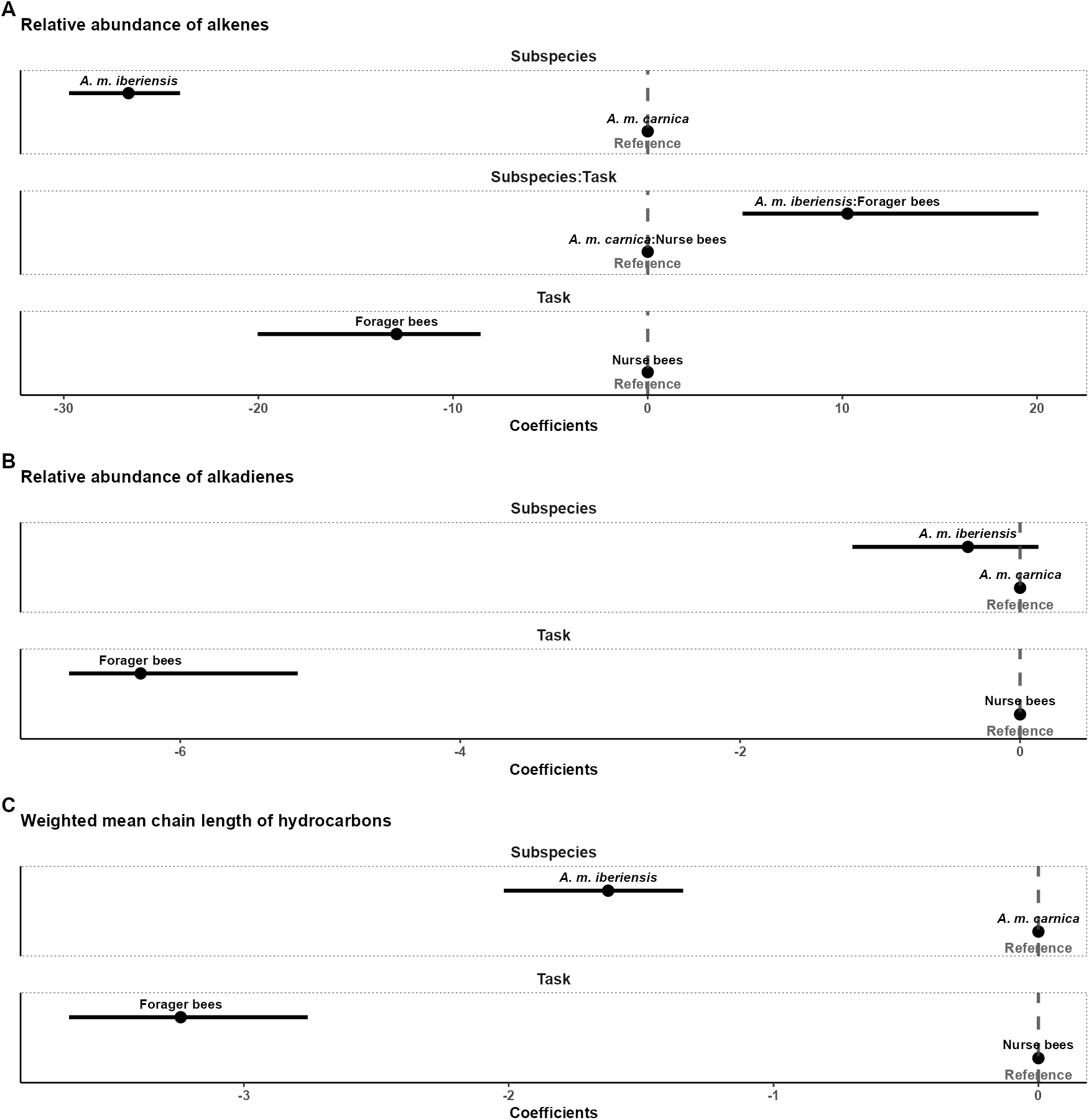
Quantile regression coefficients (effect sizes) for the task and subspecies-related differences in the CHC composition of honey bee workers. The figure is divided into three plots, each corresponding to the results of a quantile regression (50% quantile) analysis on the task- and subspecies-related differences in a compositional trait of the CHC profile of honey bee workers. **A)** Relative abundance of monounsaturated hydrocarbons (alkenes). **B)** Relative abundance of di-unsaturated hydrocarbons (alkadienes). **C)** Mean chain length of the CHCs. Each plot is divided into facets, which correspond to the categorical variables used as predictors for the model (task and subspecies) and their interaction (task:subspecies), if included. Point intervals represent the estimated difference to the reference group (task: nurse bees; subspecies: *A. m. carnica*; subspecies:task: *A. m. carnica*:nurse bees) and its 95% confidence intervals. The reference is visualized with a dashed grey line and an overlapping point for the corresponding group.

**Figure S2:**
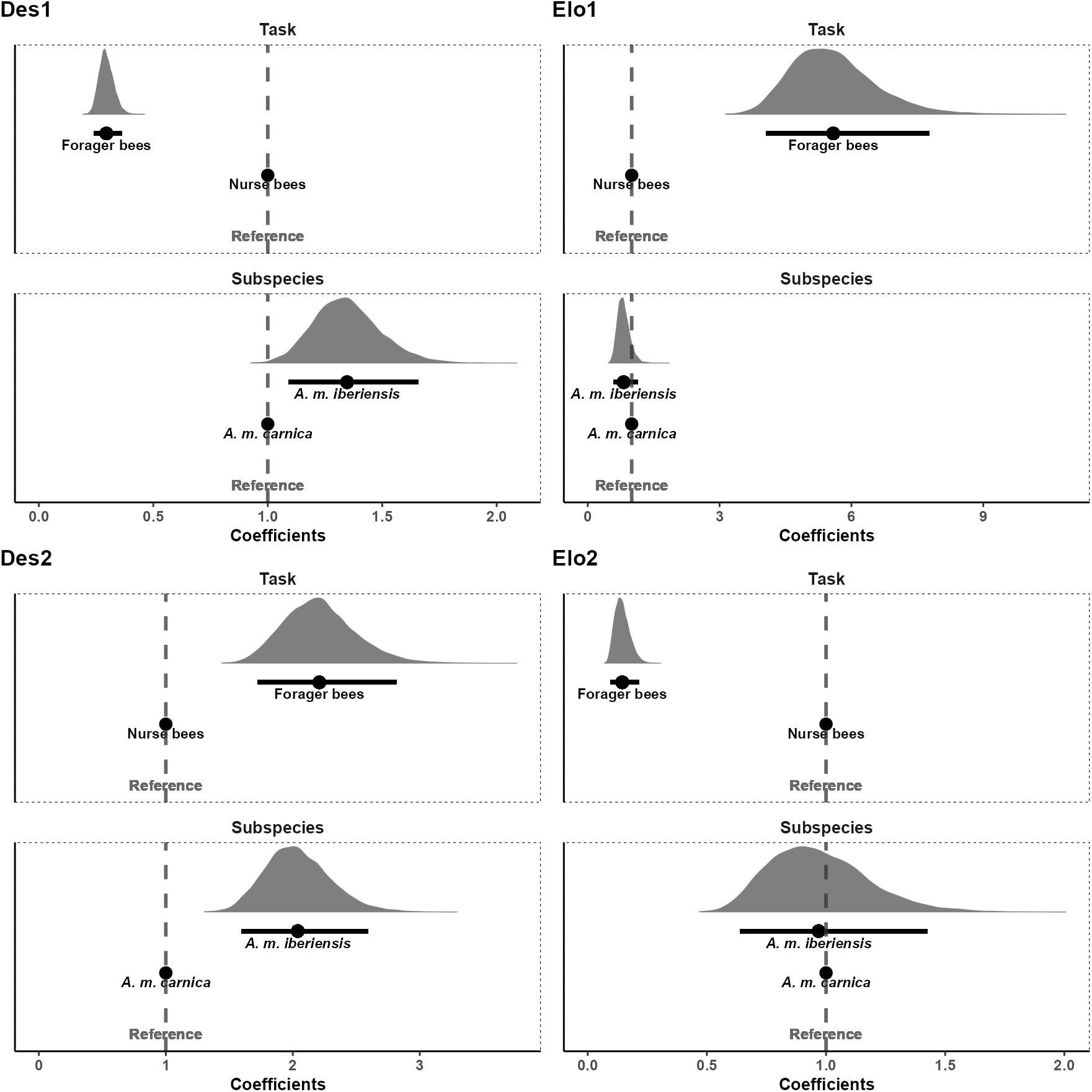
Bootstrapped GLM coefficients (effect sizes) for the difference in the relative expression of the CHCs biosynthetic genes between tasks and subspecies. The figure is divided into four plots, each corresponding to a gene. Each plot is divided into two facets, which correspond to the two categorical variables used as predictors for the model (task and subspecies). Shaded curves depict the distribution of the estimated GLM coefficients across the 10,000 bootstrap simulations. Point intervals represent the median of the bootstrapped GLM mean proportional difference in the gene expression to the reference group (task: nurse bees; subspecies: *A. m. carnica*) and its 95% confidence intervals. The reference is visualized with a dashed grey line and an overlapping point for the corresponding group.

## References

Blomquist GJ, Bagnères A-G. 2010. Insect Hydrocarbons: Biology, Biochemistry, and Chemical Ecology. Cambridge University Press

Blomquist GJ, Nelson DR, De Renobales M. 1987. Chemistry, biochemistry, and physiology of insect cuticular lipids. Archives of Insect Biochemistry and Physiology [Internet] 6:227–265. Available from: https://onlinelibrary.wiley.com/doi/abs/10.1002/arch.940060404 http://doi.wiley.com/10.1002/arch.940060404

Carlson DA, Roan CShyan, Yost RA, Hector Julio. 1989. Dimethyl disulfide derivatives of long chain alkenes, alkadienes, and alkatrienes for gas chromatography/mass spectrometry. Anal. Chem. [Internet] 61:1564–1571. Available from: 10.1021/ac00189a019

Chávez-Galarza J, Henriques D, Johnston JS, Carneiro M, Rufino J, Patton JC, Pinto MA. 2015. Revisiting the Iberian honey bee (Apis mellifera iberiensis) contact zone: Maternal and genome-wide nuclear variations provide support for secondary contact from historical refugia. Molecular Ecology [Internet] 24:2973–2992. Available from: https://onlinelibrary.wiley.com/doi/abs/10.1111/mec.13223

Chung H, Carroll SB. 2015. Wax, sex and the origin of species: Dual roles of insect cuticular hydrocarbons in adaptation and mating. BioEssays [Internet] 37:822–830. Available from: https://onlinelibrary.wiley.com/doi/abs/10.1002/bies.201500014

Cinti DL, Cook L, Nagi MN, Suneja SK. 1992. The fatty acid chain elongation system of mammalian endoplasmic reticulum. Progress in Lipid Research [Internet] 31:1–51. Available from: https://www.sciencedirect.com/science/article/pii/016378279290014A

Dallerac R, Labeur C, Jallon J-M, Knipple DC, Roelofs WL, Wicker-Thomas C. 2000. A $\Delta$9 desaturase gene with a different substrate specificity is responsible for the cuticular diene hydrocarbon polymorphism in Drosophila melanogaster. Proceedings of the National Academy of Sciences [Internet] 97:9449–9454. Available from: https://www.pnas.org/doi/full/10.1073/pnas.150243997

De la Rúa P, Jaffé R, Dall’Olio R, Muñoz I, Serrano J. 2009. Biodiversity, conservation and current threats to European honeybees. Apidologie [Internet] 40:263–284. Available from: http://link.springer.com/10.1051/apido/2009027

Denic V, Weissman JS. 2007. A Molecular Caliper Mechanism for Determining Very Long-Chain Fatty Acid Length. Cell [Internet] 130:663–677. Available from: https://www.sciencedirect.com/science/article/pii/S009286740700829X

Dogantzis KA, Tiwari T, Conflitti IM, Dey A, Patch HM, Muli EM, Garnery L, Whitfield CW, Stolle E, Alqarni AS, et al. 2021. Thrice out of Asia and the adaptive radiation of the western honey bee. Science Advances [Internet] 7:eabj2151. Available from: https://www.science.org/doi/full/10.1126/sciadv.abj2151

Falcón T, Ferreira-Caliman MJ, Franco Nunes FM, Tanaka ÉD, do Nascimento FS, Gentile Bitondi MM. 2014. Exoskeleton formation in Apis mellifera: Cuticular hydrocarbons profiles and expression of desaturase and elongase genes during pupal and adult development. Insect Biochemistry and Molecular Biology [Internet] 50:68–81. Available from: https://www.sciencedirect.com/science/article/pii/S0965174814000769

Garnery L, Cornuet J-M, Solignac M. 1992. Evolutionary history of the honey bee Apis mellifera inferred from mitochondrial DNA analysis. Molecular Ecology [Internet] 1:145–154. Available from: http://doi.wiley.com/10.1111/j.1365-294X.1992.tb00170.x

Gibbs A. 1995. Physical properties of insect cuticular hydrocarbons: Model mixtures and lipid interactions. Comparative Biochemistry and Physiology Part B: Biochemistry and Molecular Biology [Internet] 112:667–672. Available from: https://www.sciencedirect.com/science/article/pii/0305049195001190?via%3Dihub

Gibbs AG. 1998. Water-proofing properties of cuticular lipids. American Zoologist [Internet] 38:471–482. Available from: https://academic.oup.com/icb/article-lookup/doi/10.1093/icb/38.3.471

Gibbs AG. 2002. Lipid melting and cuticular permeability: New insights into an old problem. Journal of Insect Physiology [Internet] 48:391–400. Available from: https://www.sciencedirect.com/science/article/pii/S00221 https://linkinghub.elsevier.com/retrieve/pii/S0022191002000598

Gibbs A, Pomonis JG. 1995. Physical properties of insect cuticular hydrocarbons: The effects of chain length, methyl-branching and unsaturation. Comparative Biochemistry and Physiology Part B: Biochemistry and Molecular Biology [Internet] 112:243–249. Available from: https://www.sciencedirect.com/science/article/pii/030504919500081X?via%3Dihub

Greene MJ, Gordon DM. 2003. Cuticular hydrocarbons inform task decisions. Nature [Internet] 423:32–32. Available from: https://www.nature.com/articles/423032a

Gu P, Welch WH, Guo L, Schegg KM, Blomquist GJ. 1997. Characterization of a Novel Microsomal Fatty Acid Synthetase (FAS) Compared to a Cytosolic FAS in the Housefly, Musca domestica. Comparative Bio-chemistry and Physiology Part B: Biochemistry and Molecular Biology [Internet] 118:447–456. Available from: https://www.sciencedirect.com/science/article/pii/S0305049197001120

Han F, Wallberg A, Webster MT. 2012. From where did the Western honeybee (Apis mellifera) originate? Ecology and Evolution [Internet] 2:1949–1957. Available from: https://www.ncbi.nlm.nih.gov/pmc/articles/PMC3431949.pdf http://doi.wiley.com/10.1002/ece3.312

Hellemans J, Mortier G, De Paepe A, Speleman F, Vandesompele J. 2007. qBase relative quantification framework and software for management and automated analysis of real-time quantitative PCR data. Genome Biology [Internet] 8:R19. Available from: 10.1186/gb-2007-8-2-r19

Holze H, Schrader L, Buellesbach J. 2021. Advances in deciphering the genetic basis of insect cuticular hydrocarbon biosynthesis and variation. Heredity [Internet] 126:219–234. Available from: https://www.nature.com/articles/s41437-020-00380-y

Howard RW, Blomquist GJ. 2005. Ecological, behavioral, and biochemical aspects of insect hydrocarbons. Annual Review of Entomology [Internet] 50:371–393. Available from: http://www.annualreviews.org/doi/10.1146/annurev.ento.50.071803.130359

Huang Z-Y, Robinson GE. 1996. Regulation of honey bee division of labor by colony age demography. Behav Ecol Sociobiol [Internet] 39:147–158. Available from: 10.1007/s002650050276

Huang ZY, Robinson GE. 1992. Honeybee colony integration: Worker-worker interactions mediate hormonally regulated plasticity in division of labor. Proceedings of the National Academy of Sciences [Internet] 89:11726–11729. Available from: https://www.pnas.org/doi/abs/10.1073/pnas.89.24.11726

Hunt BG, Ometto L, Keller L, Goodisman MAD. 2013. Evolution at Two Levels in Fire Ants: The Relationship between Patterns of Gene Expression and Protein Sequence Evolution. Molecular Biology and Evolution [Internet] 30:263–271. Available from: 10.1093/molbev/mss234

Hunt BG, Wyder S, Elango N, Werren JH, Zdobnov EM, Yi SV, Goodisman MAD. 2010. Sociality Is Linked to Rates of Protein Evolution in a Highly Social Insect. Molecular Biology and Evolution [Internet] 27:497–500. Available from: https://academic.oup.com/mbe/article-lookup/doi/10.1093/molbev/msp225

Jakobsson A, Westerberg R, Jacobsson A. 2006. Fatty acid elongases in mammals: Their regulation and roles in metabolism. Progress in Lipid Research [Internet] 45:237–249. Available from: https://www.sciencedirect.com/science/article/pii/S0163782706000051

Johnson BR. 2008. Within-nest temporal polyethism in the honey bee. Behav Ecol Sociobiol [Internet] 62:777–784. Available from: 10.1007/s00265-007-0503-2

Johnson BR, Frost E. 2012. Individual-level patterns of division of labor in honeybees highlight flexibility in colony-level developmental mechanisms. Behav Ecol Sociobiol [Internet] 66:923–930. Available from: 10.1007/s00265-012-1341-4

Juárez P, Chase J, Blomquist GJ. 1992. A microsomal fatty acid synthetase from the integument of Blattella germanica synthesizes methyl-branched fatty acids, precursors to hydrocarbon and contact sex pheromone. Archives of Biochemistry and Biophysics [Internet] 293:333–341. Available from: https://www.sciencedirect.com/science/article/pii/000398619290403J

Kather R, Drijfhout FP, Martin SJ. 2011. Task group differences in cuticular lipids in the honey bee apis mellifera. Journal of Chemical Ecology [Internet] 37:205–212. Available from: http://link.springer.com/10.1007/s10886-011-9909-4

Koenker R. 2023. Quantreg: Quantile regression. Available from: https://CRAN.R-project.org/package=quantreg

Kuhn M, Wickham H. 2020. Tidymodels: A collection of packages for modeling and machine learning using tidyverse principles. Available from: https://www.tidymodels.org

Lenth RV. 2023. Emmeans: Estimated marginal means, aka least-squares means. Available from: https://CRAN.R-project.org/package=emmeans

M. Elekonich M, Schulz DJ, Bloch G, Robinson GE. 2001. Juvenile hormone levels in honey bee (Apis mellifera L.) foragers: Foraging experience and diurnal variation. Journal of Insect Physiology [Internet] 47:1119–1125. Available from: https://www.sciencedirect.com/science/article/pii/S0022191001000907

Makki R, Cinnamon E, Gould AP. 2014. The Development and Functions of Oenocytes. Annu Rev Entomol [Internet] 59:405–425. Available from: https://www.ncbi.nlm.nih.gov/pmc/articles/PMC7613053/

Martin SJ, Drijfhout FP. 2009. Nestmate and Task Cues are Influenced and Encoded Differently within Ant Cuticular Hydrocarbon Profiles. J Chem Ecol [Internet] 35:368–374. Available from: 10.1007/s10886-009-9612-x

Maul V, Hähnle A. 1994. Morphometric studies with pure bred stock of Apis mellifera carnica Pollmann from Hessen. Apidologie [Internet] 25:119–132. Available from: 10.1051/apido:19940201

Mayer M. 2023. Confintr: Confidence intervals. Available from: https://CRAN.R-project.org/package=confintr

Menzel Florian, Blaimer BB, Schmitt T. 2017. How do cuticular hydrocarbons evolve? Physiological constraints and climatic and biotic selection pressures act on a complex functional trait. Proceedings of the Royal Society B: Biological Sciences [Internet] 284:20161727. Available from: 10.1098/rspb.2016.1727 http://rspb.royalsocietypublishing.org/lookup/doi/10.1098/rspb.2016.17

Menzel F, Morsbach S, Martens JH, Räder P, Hadjaje S, Poizat M, Abou B. 2019. Communication versus waterproofing: The physics of insect cuticular hydrocarbons. The Journal of Experimental Biology [Internet] 222:jeb210807. Available from: http://www.ncbi.nlm.nih.gov/pubmed/31704903 http://jeb.biologists.org/lookup/doi/10.1242/jeb.210807

Menzel F, Schmitt T, Blaimer BB. 2017. The evolution of a complex trait: Cuticular hydrocarbons in ants evolve independent from phylogenetic constraints. Journal of Evolutionary Biology [Internet] 30:1372–1385. Available from: http://doi.wiley.com/10.1111/jeb.13115

Menzel F, Zumbusch M, Feldmeyer B. 2018. How ants acclimate: Impact of climatic conditions on the cuticular hydrocarbon profile. Functional Ecology 32:657–666.

Moris VC, Podsiadlowski L, Martin S, Oeyen JP, Donath A, Petersen M, Wilbrandt J, Misof B, Liedtke D, Thamm M, et al. 2023. Intrasexual cuticular hydrocarbon dimorphism in a wasp sheds light on hydrocarbon biosynthesis genes in Hymenoptera. Commun Biol [Internet] 6:1–15. Available from: https://www.nature.com/articles/s42003-022-04370-0

Oksanen J, Simpson GL, Blanchet FG, Kindt R, Legendre P, Minchin PR, O’Hara RB, Solymos P, Stevens MHH, Szoecs E, et al. 2022. Vegan: Community ecology package. Available from: https://CRAN.R-project.org/package=vegan

Ottensmann M, Stoffel MA, Nichols HJ, Hoffman JI. 2018. GCalignR: An R package for aligning gaschromatography data for ecological and evolutionary studies. PLOS ONE.

Pedersen TL. 2023. Patchwork: The composer of plots. Available from: https://CRAN.R-project.org/package=patchwork

Pedersen TL, Shemanarev M. 2023. Ragg: Graphic devices based on AGG. Available from: https://CRAN.R-project.org/package=ragg

Posit Team. 2024. RStudio: Integrated development environment for R. Boston, MA Available from: http://www.posit.com/

Qiu Y, Tittiger C, Wicker-Thomas C, Le Goff G, Young S, Wajnberg E, Fricaux T, Taquet N, Blomquist GJ, Feyereisen R. 2012. An insect-specific P450 oxidative decarbonylase for cuticular hydrocarbon biosynthesis. Proceedings of the National Academy of Sciences [Internet] 109:14858–14863. Available from: https://www.pnas.org/doi/abs/10.1073/pnas.1208650109

R Core Team. 2024. R: A language and environment for statistical computing. Vienna, Austria Available from: https://www.R-project.org/

Reed JR, Quilici DR, Blomquist GJ, Reitz RC. 1995. Proposed mechanism for the cytochrome P 450-catalyzed conversion of aldehydes to hydrocarbons in the house fly, Musca domestica. Biochemistry [Internet] 34:16221–16227. Available from: https://pubs.acs.org/doi/abs/10.1021/bi00049a038

Reim T, Scheiner R. 2014. Division of labour in honey bees: Age- and task-related changes in the expression of octopamine receptor genes. Insect Molecular Biology [Internet] 23:833–841. Available from: https://onlinelibrary.wiley.com/doi/abs/10.1111/imb.12130

Robinson GE, Page Jr. RE, Strambi C, Strambi A. 1992. Colony Integration in Honey Bees: Mechanisms of Behavioral Reversion. Ethology [Internet] 90:336–348. Available from: https://onlinelibrary.wiley.com/doi/abs/10.1111/j.1439-0310.1992.tb00844.x

Rodríguez-Leon DS. 2023b. easyqpcr2: Custom package based on the abandoned EasyqpcR package.

Rodríguez-Leon DS. 2023a. analyzeGC: Analysis of GC data. Available from: https://github.com/dsrodriguezl/analyzeGC

Rodríguez-León DS, Uzunov A, Costa C, Dylan ES, Charistos L, Galea T, Gabel M, Scheiner R, Pinto MA, Schmitt T. 2024. Deciphering the variation in cuticular hydrocarbon profiles of six European honey bee subspecies. bioRxiv [Internet]:2024.07.05.601031. Available from: http://biorxiv.org/content/early/2024/07/20/2024.07.05.601031.abstract

Ruttner F. 1988. Biogeography and taxonomy of honeybees. Berlin, Heidelberg: Springer Berlin Heidelberg Available from: https://link.springer.com/book/10.1007/978-3-642-72649-1

Ruttner F, Tassencourt L, Louveaux J. 1978. BIOMETRICAL-STATISTICAL ANALYSIS OF THE GEOGRAPHIC VARIABILITY OF APIS MELLIFERA L.* I. Material and Methods BIOMETRICAL-STATISTICAL ANALYSIS OF THE GEOGRAPHIC VARIABILITY OF APIS MELLIFERA L.* I. Material and Methods Institutfi7r Bienenkunde, D-6370 Ober. Springer Verlag Available from: https://hal.archives-ouvertes.fr/hal-00890475

Schal C, Sevala VL, Young HP, Bachmann JAS. 1998. Sites of Synthesis and Transport Pathways of Insect Hydrocarbons: Cuticle and Ovary as Target Tissues1. American Zoologist [Internet] 38:382–393. Available from: 10.1093/icb/38.2.382

Scheiner R, Reim T, Søvik E, Entler BV, Barron AB, Thamm M. 2017. Learning, gustatory responsiveness and tyramine differences across nurse and forager honeybees. Journal of Experimental Biology [Internet] 220:1443–1450. Available from: 10.1242/jeb.152496

Scheiner R, Toteva A, Reim T, Søvik E, Barron AB. 2014. Differences in the phototaxis of pollen and nectar foraging honey bees are related to their octopamine brain titers. Front. Physiol. [Internet] 5. Available from: https://www.frontiersin.org/journals/physiology/articles/10.3389/fphys.2014.00116/full

Schilcher F, Scheiner R. 2023. New insight into molecular mechanisms underlying division of labor in honeybees. Current Opinion in Insect Science [Internet]:101080. Available from: https://www.sciencedirect.com/science/article/pii/S2214574523000779

Siegel AJ, Fondrk MK, Amdam GV, Page RE. 2013. In-hive patterns of temporal polyethism in strains of honey bees (Apis mellifera) with distinct genetic backgrounds. Behav Ecol Sociobiol [Internet] 67:1623–1632. Available from: 10.1007/s00265-013-1573-y

Smith CR, Toth AL, Suarez AV, Robinson GE. 2008. Genetic and genomic analyses of the division of labour in insect societies. Nat Rev Genet [Internet] 9:735–748. Available from: https://www.nature.com/articles/nrg2429

Takahashi A, Tsaur S-C, Coyne JA, Wu C-I. 2001. The nucleotide changes governing cuticular hydrocarbon variation and their evolution in Drosophila melanogaster. Proceedings of the National Academy of Sciences [Internet] 98:3920–3925. Available from: https://www.pnas.org/doi/full/10.1073/pnas.061465098

Tiedemann F. 2022. Gghalves: Compose half-half plots using your favourite geoms. Available from: https://CRAN.R-project.org/package=gghalves

Vernier CL, Krupp JJ, Marcus K, Hefetz A, Levine JD, Ben-Shahar Y. 2019. The cuticular hydrocarbon profiles of honey bee workers develop via a socially-modulated innate process. eLife [Internet] 8. Available from: https://elifesciences.org/articles/41855

Wagner D, Brown MJF, Broun P, Cuevas W, Moses LE, Chao DL, Gordon DM. 1998. Task-Related Differences in the Cuticular Hydrocarbon Composition of Harvester Ants, Pogonomyrmex barbatus. Journal of Chemical Ecology [Internet] 24:2021–2037. Available from: http://link.springer.com/10.1023/A:1020781508889

Wagner D, Tissot M, Gordon D. 2001. Task-related environment alters the cuticular hydrocarbon composition of harvester ants. Journal of Chemical Ecology [Internet] 27:1805–1819. Available from: http://link.springer.com/10.1023/A:1010408725464

Wallberg A, Han F, Wellhagen G, Dahle B, Kawata M, Haddad N, Simões ZLP, Allsopp MH, Kandemir I, De la Rúa P, et al. 2014. A worldwide survey of genome sequence variation provides insight into the evolutionary history of the honeybee Apis mellifera. Nat Genet [Internet] 46:1081–1088. Available from: https://www.nature.com/articles/ng.3077%3E%C2%A0

Wickham H, Averick M, Bryan J, Chang W, McGowan LD, François R, Grolemund G, Hayes A, Henry L, Hester J, et al. 2019. Welcome to the tidyverse. Journal of Open Source Software 4:1686.

Wigglesworth VB. 1933. Memoirs: The Physiology of the Cuticle and of Ecdysis in Rhodnius prolixus (Triatomidae, Hemiptera); with special reference to the function of the oenocytes and of the dermal glands. Journal of Cell Science [Internet] s2-76:269–318. Available from: 10.1242/jcs.s2-76.302.269

Wilke CO, Wiernik BM. 2022. Ggtext: Improved text rendering support for ‘ggplot2’. Available from: https://CRAN.R-project.org/package=ggtext

Winston ML, Fergusson LA. 1985. The effect of worker loss on temporal caste structure in colonies of the honeybee (Apis mellifera L.). Can. J. Zool. [Internet] 63:777–780. Available from: https://cdnsciencepub.com/doi/abs/10.1139/z85-113

